## Supplementary material for "UGGT1-mediated reglucosylation of *N*-glycan competes with ER-associated degradation of unstable and misfolded glycoproteins": Key resources Table-1

| Reagent type or resources | Designation | Source or reference | Identifier | Additional information |
|---|---|---|---|---|
| Cell line (*Homo sapiens* ) | colorectal carcinoma | ATCC | HCT116 | Parental HCT116 cell line have been authenticate |
| Recombinant DNA reagent | p3xFlag-CMV-14 | Ninagawa et al., 2014 | | |
| Recombinant DNA reagent | DT-A-pA/loxP/PGK-Puro-pA/loxP | Ninagawa et al., 2014 | | |
| Recombinant DNA reagent | DT-A-pA/loxP/PGK-Hygro-pA/loxP | Tsuda et al., 2019 | | |
| Antibody | anti-HA (Rabbit polyclonal) | Recenttec | Cat#: R4-TP1411100 | WB (1:1000) |
| Antibody | anti-HA (mouse polyclonal) | Recenttec | Cat#: R4-TM1422100 | WB (1:1000) |
| Antibody | anti-UGGT1 (Rabbit polyclonal) | Sigma | Cat#: HPA015127 | WB (1:1000) |
| Antibody | anti-UGGT2 (Rabbit polyclonal) | GeneTex | Cat#: GTX103837 | WB (1:1000) |
| Antibody | anti-GAPDH (Rabbit polyclonal) | Trevigen | Cat#: 2275-PC-100 | WB (1:1000) |
| Antibody | anti-Myc (Rabbit polyclonal) | MBL | Cat#: MBL562 | WB (1:1000) |
| Antibody | anti-Myc (Mouse monoclonal) | Wako | Cat#: 011-21874 | WB (1:1000) |
| Antibody | anti-ATF6 (Rabbit polyclonal) | Haze et al., 1999 | | WB (1:4000) |
| Antibody | anti-A1AT (Rabbit polyclonal) | Dako | Cat#: A0012 | WB (1:1000) |
| Antibody | anti-Flag (Mouse monoclonal) | Sigma | Cat#: F3165 | WB (1:1000) |
| Antibody | anti-Grp170 (Rabbit polyclonal) | GeneTex | Cat#: 102255 | WB (1:1000) |
| Antibody | anti-Sil1 (Rabbit polyclonal) | GeneTex | Cat#: GTX116755 | WB (1:1000) |
| Antibody | anti-Ribophorin I (Rabbit monoclonal) | abcam | Cat#: ab198508 | WB (1:1000) |
| Antibody | anti-CRT (Rabbit monoclonal) | Enzo Life Sciences | Cat#: ADI-SPA-600 | WB (1:1000) |
