## Supplementary material for "UGGT1-mediated reglucosylation of *N*-glycan competes with ER-associated degradation of unstable and misfolded glycoproteins": Used primers

**Supplementary File 1**

| Reagent type or resource | Designation | Source or reference | Identifier | Additional information |
|---|---|---|---|---|
| Sequence-based reagent | UGGT1_PCRcheckFw | This paper | Genomic PCR primer | GTCATTGGGTTAGTGCCCAC |
| Sequence-based reagent | UGGT1_PCRcheckRv | This paper | Genomic PCR primer | GTGCCCAGCCTTCTTCCACG |
| Sequence-based reagent | UGGT2_PCRcheckFw | This paper | Genomic PCR primer | TGGCCCTGGAATGACATTAC |
| Sequence-based reagent | UGGT2_PCRcheckRv | This paper | Genomic PCR primer | TACAATTGACTAGGTACCTGAG |
| Sequence-based reagent | SEL1L_PCRcheckFw | This paper | Genomic PCR primer | AAGGGCAGGCACACCAAGTC |
| Sequence-based reagent | SEL1L_PCRcheckRv | This paper | Genomic PCR primer | TGACAGGTCACCGCTTTCTCA |
| Sequence-based reagent | UGGT1-PuroFw | This paper | Primer for vector PCR | GGGAGTTCTGGTTGTACTCACTGTTGAACCTCTTCGAGGGACCTA |
| Sequence-based reagent | UGGT1-PuroRv | This paper | Primer for vector PCR | CCTTTACTGAGGAGAACAGCCACAGCATATTCAATAACCCTTAAT |
| Sequence-based reagent | UGGT1-BackboneFw | This paper | Primer for vector PCR | GGAAAGAGGACATTTTTCCCTTAAGCCACAGAACAGTGAGTACAACCATAGGTCCCTCGAAGAGGTTCACTAG |
| Sequence-based reagent | UGGT1-BackboneRv | This paper | Primer for vector PCR | GTAATGTCAACTTATTTGTAAATACCCACAGAACAGTGAGTACAACCAATTAAGGGTTATTGAATATGATCGG |

| | | | | |
|---|---|---|---|---|
| Sequence-based reagen | UGGT1-LarmFw | This paper | Genomic PCR primer | GTATTTACAAATAAGTTGAC |
| Sequence-based reagen | UGGT1-LarmRv | This paper | Genomic PCR primer | AACAGTGAGTACAACCAGAAC |
| Sequence-based reagent | UGGT1-RarmFw | This paper | Genomic PCR primer | CTGTGGCTGTTCTCCTCAGTAAAG |
| Sequence-based reagent | UGGT1-RarmRv | This paper | Genomic PCR primer | CTTAAGGGAAAAATGTCCTC |
| Sequence-based reagent | UGGT1-sgRNAFw | This paper | Primer for vector PCR | GTTGTACTCACTGTTCTGgttttagagctaGAAAtagc |
| Sequence-based reagent | UGGT1-sgRNARv | This paper | Primer for vector PCR | GGTGTTTCGTCCTTTCCAC |
| Sequence-based reagent | UGGT2-HygroFw | This paper | Primer for vector PCR | GATTGCAGCTGATGAGCCACCACCAGAACCTCTTCGAGGGACCTA |
| Sequence-based reagent | UGGT2-HygroRv | This paper | Primer for vector PCR | TAACCACAAATGCATTACAACCATCCATATTCAATAACCCTTAAT |
| Sequence-based reagent | UGGT2-BackboneFw | This paper | Primer for vector PCR | TTTACTAATTCTTTCTGTAGCTTTCCCACCAGATGGTTGTAATGCATTACTAGTTCTAGAGCATTTAAATACG |
| Sequence-based reagent | UGGT2-BackboneRv | This paper | Primer for vector PCR | TTTGGATCTTCCAGTAATGTCCAAGCCACCAGATGGTTGTAATGCATTATCGGAATTCGATAGCGGCCGCTGG |
| Sequence-based reagent | UGGT2-LarmFw | This paper | qRT-PCR primer | CTTGGACATTACTGGAAGATC |
| Sequence-based reagent | UGGT2-LarmRv | This paper | qRT-PCR primer | TGGTGGTGGCTCATCAGCTGC |
| Sequence-based reagent | UGGT2-RarmFw | This paper | qRT-PCR primer | GATGGTTGTAATGCATTTGTG |
| Sequence-based reagent | UGGT2-RarmRv | This paper | qRT-PCR primer | GAAAGCTACAGAAAGAATTAG |

| | | | | |
|---|---|---|---|---|
| Sequence-based reagent | UGGT2-sgRNAFw | This paper | Primer for vector PCR | AATGCATTACAACCATCTGGgttttagagctaGAAAtagc |
| Sequence-based reagent | UGGT2-sgRNARv | This paper | Primer for vector PCR | GGTGTTTCGTCCTTTCCAC |
| Sequence-based reagent | UGGT1-cloningFw | This paper | RT-PCR primer | ATAAGAATGCGGCCGCggcatgggctgcaagggagac |
| Sequence-based reagent | UGGT1-cloningRv | This paper | RT-PCR primer | gcTCTAGAtaattcttcacgtttctgag |
| Sequence-based reagent | UGGT2-cloningFw | This paper | RT-PCR primer | ATAAGAATGCGGCCGCgccatggcgccagcgaaagc |
| Sequence-based reagent | UGGT2-cloningRv | This paper | RT-PCR primer | ggGGTACCgaGAGTTCATCATGTGTCAAAATTG |
| Sequence-based reagent | UGGT1-D1358A-Fw | This paper | Primer for vector | TTTGTGGCAGCTGATCAGATTGTACGAACAGATCT |
| Sequence-based reagent | UGGT1D1358A-Rv | This paper | Primer for vector | AATCTGTCAGCTGCCACAACAGGAACTTGTCAAC |
| Sequence-based reagent | SEL1L-clon-Fw | Ninagawa et al., 2011 CSF{Ninagawa, 2011 #263} | RT-PCR primer | CCATCGATAGGATGCGGGTCCGGATAGGGC |
| Sequence-based reagent | SEL1L-clon-Rv | Ninagawa et al., 2011 CSF {Ninaga | RT-PCR primer | CGGGATCCCTGTGGTGGCTGCTGCTCTG |

| | | | | |
|---|---|---|---|---|
| | | wa, 2011 #263} | | |
| Sequence-based reagent | 3xMyc-Fw | This paper | Primer for vector | CGGGATCCTGTGGTGGAGTTCTGGAGC |
| Sequence-based reagent | 3xMyc-Rv | This paper | Primer for vector | CGGGATCCAAATTCTCTCAAGACAGGTC |
| Sequence-based reagent | RatRI332-cloningFw | This paper | RT-PCR primer | CCCAAGCTTGCGGTCATGGAGGCGCCGATCGTCTT |
| Sequence-based reagent | RatRI332-cloningRv | This paper | RT-PCR primer | GGGGTACCCCTACAAACCGCATCTTCAGTG |
